# Expansion, Exploitation and Extinction: Niche Construction in Ephemeral Landscapes

**DOI:** 10.1101/489096

**Authors:** Miles T. Wetherington, Juan E. Keymer

**Affiliations:** Department of Ecology, School of Biological Sciences, P. Catholic University of Chile, Santiago, Chile; Institute of Physics, School of Physics, P. Catholic University of Chile, Santiago, Chile; Facultad de Ciencias Biológicas, Pontificia Universidad Católica de Chile, Avda. Libertador Bernardo O’Higgins 340, Santiago, Chile; Instituto de Física Pontificia Universidad Católica de Chile, Avda. Vicuña Mackenna 4860, Santiago, Chile

**Author notes:** Both authors developed the model, MTW analyzed the data, and both authors wrote the manuscript. MTW (Miles T Wetherington) and JEK (Juan E Keymer) contributed equally to this work.

**Keywords:** Niche Construction, Metapopulation Dynamics, Critical Transitions, Ecological Suicide, Ephemeral Landscapes

## Abstract

We developed an interacting particle system (IPS) to study the effect of niche construction on metapopulation dynamics in ephemeral landscapes. Using finite scaling theory, we find a divergence in the qualitative behavior at the extinction threshold between analytic (mean field) and numerical (IPS) results when niche construction is confined to a small area in the spatial model. While increasing the area of niche construction leads to a faster rate of range expansion, it also causes a shift from a continuous to discontinuous phase transition, thus returning to the mean field prediction. Furthermore, in the discontinuous regime of the IPS, spatial clustering prior to a critical transition disappears. This is a significant finding as spatial clustering has been considered to be an early warning signal before ecosystems reach their ‘tipping point’. In addition to maintaining stability, we find this local niche construction strategy has an advantage when in competition with an exploiter because of their ability to monopolize the constructed niche due to spatial adjacency. However, as the niche construction neighborhood expands this advantage disappears and the exploiter strategy out-competes the niche constructor. In some cases the exploiter pushes the niche constructor to extinction, thus a tragedy of the commons ensues leading to ecological suicide and a collapse of the niche.

**Significance Statement:** Many populations do not exist passively in their environment. Niche construction theory tells us the co-regulatory feedback between a population and its localized environment can have significant impacts on the evolutionary trajectory and ecological stability of the population and broader community. However, if this feedback is not tightly coupled to the niche construction strategy it is left susceptible to exploitation. In some instances this can lead to ecological suicide, where both constructor and exploiter are driven to extinction. This study emphasizes the importance of being spatial when considering the ecological implications of niche construction. Along with adding to the development of niche construction theory, this work has implications for our understanding of critical transitions in vulnerable ecosystems.

---

Understanding the co-regulatory feedback between living systems and their environment is a primary goal driving ecological research (1–3). In principle, this entails the emergence and/or maintenance of self, whether that self is a single cell (4), multicellular organism (5), population (6) or the biosphere (7). Due to the breadth of this concept, a primary task has been to partition it into spatial and temporal scales (8), as well as identify the critical units of selection (9, 10). Over the past quarter-century, research has primarily focused on studying the differences between ecological and evolutionary dynamics (3, 11), most notable of these are the closely related concepts, ecosystem engineering and niche construction. Ecosystem engineering occurs within the lifetime of the individual in a population, and is defined as its modification of the microenvironment within which it makes a living. In turn, the consequences of these actions have an impact on the coupled physical environment and ecological community (i.e. the ecotope (12)). Ecosystem engineers play an important role in the restoration and maintenance of the landscape (13), the resilience of ecosystems (14) and are often regarded as keystone species due to their role in stabilizing communities (11, 15).

An ecological interpretation of niche construction includes the mechanisms of ecosystem engineering along with all byproducts comprising the life-history of the organism (behavior, nutrient uptake, and excretion for example detritus (16), etc.) which may be considered more indirect forms of modification (17). Furthermore, niche construction has a tighter feedback to the population in question and can play an important role in influencing their evolutionary trajectory. In this way niche construction is a life history trait which has the potential to expand or maintain a populations niche, including the trans-generational inheritance of improved local conditions (ecological inheritance). The outcome of ecological inheritance for the niche constructor strategy is of particular interest for its potential to lead to further niche expansion and potentially the division of labor and differentiation within the population (5, 18). Alternatively, once this strategy emerges in the biosphere, it becomes vulnerable to exploitation, either from within the population or nearby strategies in competition for similar resources. If exploitation does not eliminate niche construction and this new exploiter-victim interaction becomes tightly coupled, exploiters will continue to profit by expanding into otherwise inaccessible regions (19). Investigating the conditions under which these outcomes are likely to occur in this way provides a mechanistic approach for understanding eco-evolutionary dynamics at the scale of the landscape (2).

The landmark work of Krakauer et al. addressed ecological and evolutionary consequences of niche construction in a Lotka-Voterra competition framework (20). We will briefly describe their system and summarize some of their findings as their work has inspired the model considered in this investigation. They introduce the constructed niche directly as an additional state variable in the dynamical system. If the niche is invested in by the developing population it acts as a public good, expanding the *niche-space* of the population. In this way the abiotic environment feeds back into the population dynamics influencing their *fundamental niche*. There is a cost to investing in the niche, notably, allocating resources (energy/time/proportion of the local population) towards its construction and maintenance (*c*), one is inherently appropriating them away from propagule production (*β*). In the case where the niche must be constantly invested in, due to external perturbations for example, a balance between production and construction must be found. This dilemma can be resolved via specialization within the population giving rise to a division of labor. Classic examples being the specialization of tasks between germ-line and somatic cells (5, 21) or the caste system in insect societies (22).

In its simplest form (logistic growth of a single strategy) the trade-off between niche construction and reproduction maximizes the *realized niche* at moderate levels of resource allocation (*c ≈ β*). However, when placed in competition with other demes the niche acts as a common-pool resource, in other words it is nonexcludable and limited. Therefore, if 2 demes compete and extract from this common-pool resource, the one with the lowest allocation towards niche construction dominates, eventually pushing the other to extinction (23). The faster producer, in this case the deme free-riding on the constructed niche, wins in the struggle for existence. Notice, however, in driving the other strategy to extinction the free-rider loses access to the portion of the niche that was constructed. Logically this leaves us with the dilemma that evolutionary and ecological dynamics should push niche construction as a strategy to extinction. Herein lies the tragedy of the commons, where the resource in question has been manifested by niche construction (24). In an environment where habitat renewal is not free and the *fundamental niche* is not inherent to the environment, for example cases where disturbances are frequent (25), local facilitation is necessary for establishment (see (26) for a review) or the ecosystem is susceptible to critical transitions (27), this scenario can lead to a form of evolutionary suicide ((28) and see (29) for a review).

With this dilemma laid out before us, how does niche construction develop and survive as a strategy in the first place? In order to reconcile biodiversity and niche construction from which it emerges (7, 30), the authors recognized the necessity to monopolize ones constructed niche, in doing so the niche constructor is less prone to exploitation by others. Therefore, the accessibility of the constructed niche can range from a completely privatized good to a common pool resource. Defining this transition from monopolized to exploited can most easily be viewed as a spatial problem.

## A. Spatial Niche Construction

Much like the insights of considering space explicitly in evolutionary games (31), a spatial instantiation of the constructed niche is a step in the right direction. As stated previously the physical environment is a necessary condition for defining the niche. This is supported by the fact that competition for space is generic to most ecosystems (32). For cases of limited dispersal, localized efforts in niche construction or a heterogeneous distribution of resources between patches, observations suggest aggregation plays an important role in the benefits of niche construction (33–37). Since it is a broad concept, we narrow our view of niche construction to the mechanism by which a population renews the necessary and sufficient conditions for future existence in its surrounding environment. Opposing this effort is the dissipation of order from the external environment, for example disturbances observed in ecosystems. Disturbances are a characteristic found at all scales (meteorite impacts, forest fires, antibiotic injections ect.) and are often a catalyst for ecological succession. Furthermore, the impact of disturbances in both landscape connectivity (38–40) and community dynamics (41, 42) are of immediate importance to the conservation of biodiversity and ecosystem services (43).

## 1. The Model

### A. The Contact Process

As a point of departure for our model, we begin by introducing the contact process (CP) (44), an interacting particle system *ξ_t_* (45) which follows colonization and extinction dynamics of particles on a countable set of spatial locations 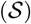 (sites). For our purposes we will consider 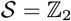, which corresponds to the set of points in 2-dimensional space with integer coordinates (lattice). On this lattice (ℤ_2_) particles (occupied sites) die out (at rate *δ*) and recruit adjacent vacant sites (at rate *β*) stochastically and in continuous time. When we say some event happens at rate *q* this signifies that given parameter *q* the time between occurrences has an exponential distribution *P*(*t_i_* ≤ *t*) = 1 − *exp*(−*qt*) and a mean of 1/*q*. The state of a site in the lattice (*x,y* ∈ ℤ_2_) can be either vacant (*ξ_t_*(*x*) = Ø) or occupied (*ξ_t_*(*x*) = 1). Denoting these states as Ø*_x_* and 1_*x*_, respectively, we are left with the defining reactions of the contact process:

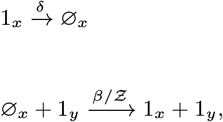

Where 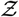 is the number of neighbors in a sites neighborhood. While an exact analytic solution to the CP is not feasible due to its spatial structure, several important features have been proven rigorously and its significance in the study of absorbing state phase transitions is well documented (46). A first attempt to understanding the dynamics of the model requires an investigation into the mean field approximation (MFA). This approach assumes the system to be ‘well-mixed’, in other words, we disregard spatial structure and assume each site is equally likely to interact with all other sites leaving us with the following evolution equation:

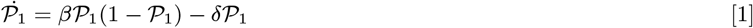

Setting 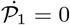 and solving for 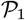 we arrive at the steady state equilibrium:

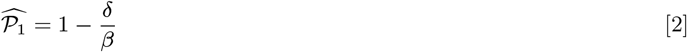

In order to maintain a viable population, propagule production must outweigh mortality (*β* > *δ*) (47). Alternatively, we can substitute in the basic reproductive number 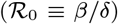 which allows us to describe the phase of the system with one parameter (48):

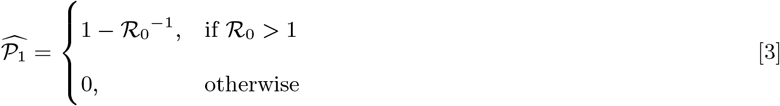

Where 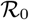 contains all the information necessary about the life history strategy and thus its long-term occupancy in the mean field limit. Setting *β* ≡ 1 we define the mean field critical mortality as *δ_c_* = 1, compared to the contact process, where *δ_c_* ≈ 0.6065. The diminished *δ_c_* in the CP is due to negative density dependence, where the crowding of particles leads to some portion of propagules to fall on already occupied sites. Spatial structure adds limitations to the efficacy of colonization, this is emphasized as we approach *δ_c_*.

### B. Ephemeral Landscapes

Next, we consider a landscape where patch lifetime is ephemeral (*sensu* Keymer et al. (39)), therefore, aside from lattice sites being occupied or vacant we add a third possible state, destroyed. This signifies a degradation in the local habitat thus leaving it unavailable for immediate colonization. All sites, regardless of state, are destroyed at rate *e*, if no mechanism for habitat renewal exists, the entire lattice converges to completely destroyed 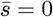. Keymer et al. investigated the extinction threshold of the contact process on a dynamic landscape defined by global rates of habitat destruction (*e*) and renewal (λ) where they envisioned λ as an ecosystem service independent of the metapopulation dynamics. Aside from the necessary conditions for the renormalized life-history of the population (i.e. 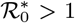), the extinction threshold is dependent on two aspects of the landscape;

i. patch lifetime,

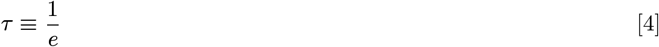
ii. and, long-term suitable habitat,

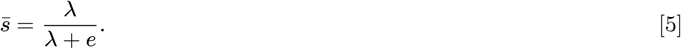

If patch lifetime is shorter than some critical time span (*τ* < *τ_min_*), the dynamic corridors generated by habitat renewal are too short lived for populations to navigate through via colonization of vacant patches, likewise, below a minimum amount of suitable habitat 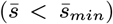, a spanning cluster of destroyed sites percolates the landscape leaving clusters of populations fragmented from suitable habitat (38). In both cases the metapopulation enters the absorbing state (extinction).

In order to connect metapopulation dynamics to the state of the landscape, we introduce a niche constructor strategy (*ξ_t_*(*x*) = 2) which, along with having the capacity to recruit local vacant sites (*ξ_t_*(*x*) = 1) for propagule production, is also capable of converting local destroyed sites (*ξ_t_*(*x*) = Ø) to vacant for future propagule production, leaving us with the following set of reactions (Figure 1 for schematic):

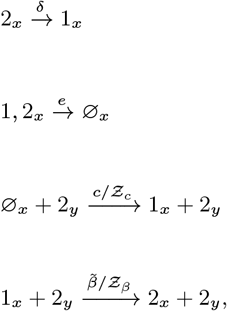

**Fig. 1.**
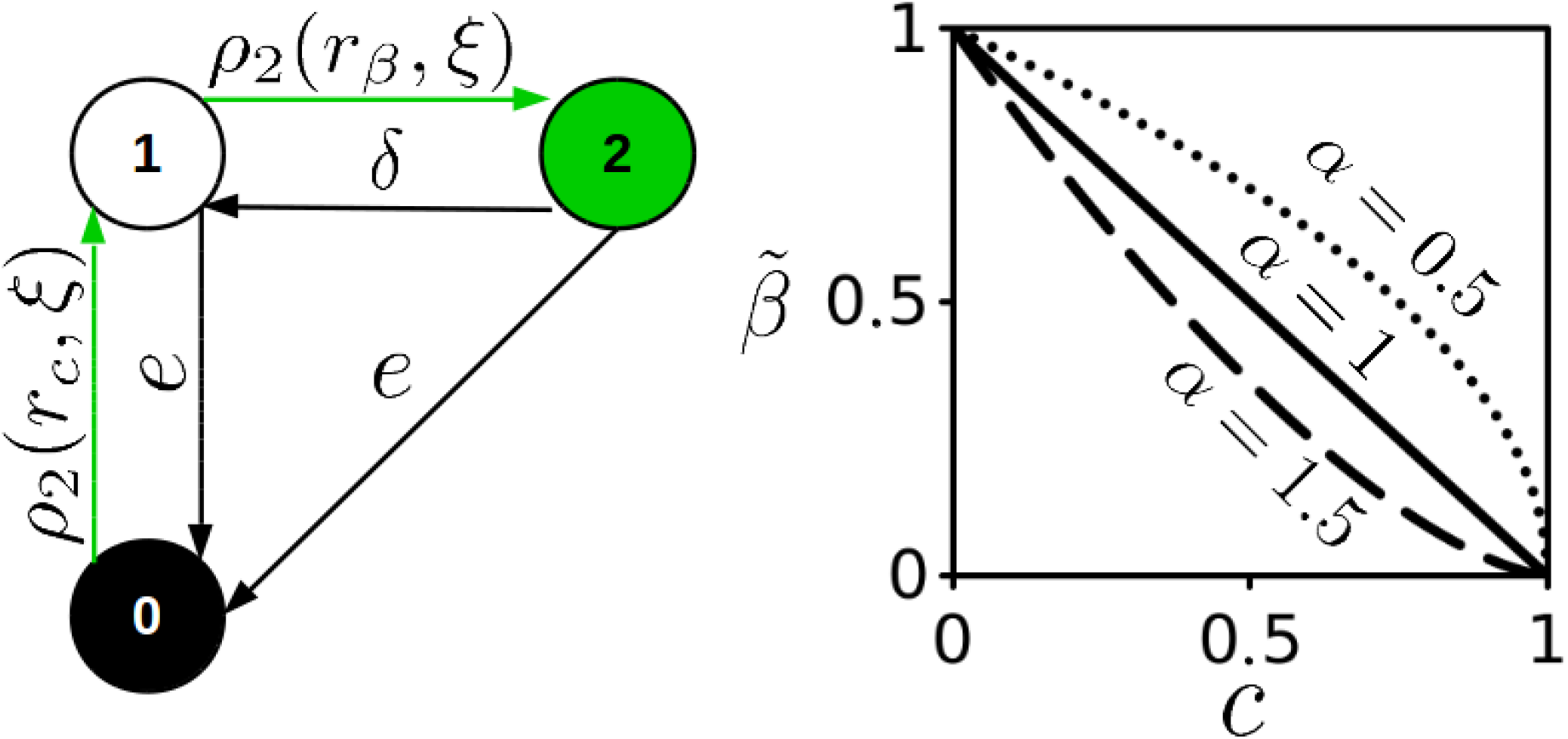
Left: Schematic showing the reactions for the niche construction model. The probability of propagule production *β* and niche construction *c* events occurring are dependent on the density (*ρ*) of occupied sites in the local neighborhood *r* of the stochastic process (*ξ*). Sites in the lattice acquire one of 3 possible states, 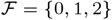 so the state of the system at time *t* is 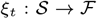. Right: Trade-off between niche construction (x-axis) and propagule production (y-axis) with respect to 3 different values of *α*.

Notice that neighborhood size 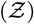 for propagule production 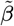 and habitat renewal *c* are not necessarily the same. Furthermore, the capacity for habitat renewal comes at an energetic cost to immediate propagule production rate:

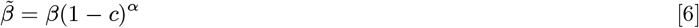

where the value of *α* determines the efficacy of niche construction.

Replacing the global ecosystem service rate *λ* with one dependent on the state of local populations we can study the co-regulation between life-history and landscape suitability.

The model described above has 4 parameters; propagule production 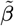, local extinction *δ*, patch destruction *e*, and patch construction *c*. The mean field approximation for this version of the model will be introduced next, followed by a comparison to the interacting particle system. After presenting these results we make an addition to the model where we add a second strategy (the basic contact process, where *c* = 0) which competes for colonizable space. We then discuss the implications of the niche construction model in the context of landscape dynamics and competition.

### C. Mean Field Approximation

Here we consider the mean field approximation for the model with the niche constructor strategy in an ephemeral landscape. Since global densities sum to unity we substitute for one of the states. For our purposes, we chose the global density of vacant patches 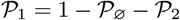. Now we can write down the mean-field approximation by considering the dynamics of destroyed and occupied patches:

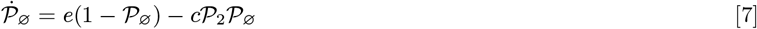

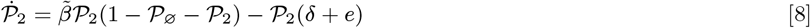

Besides the absorbing state 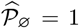 there exists a steady state equilibrium 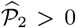. Although the exact solution is not particularly insightful (See Appendix), some observations are in order: we can determine the long-term suitable habitat

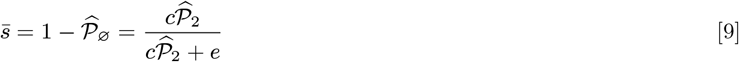

Note, 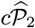 can be defined as the total constructed niche generated by the metapopulation. Substituting in the following 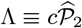 we have returned to the expression for suitable habitat (39) which is now coupled to the state of the metapopulation:

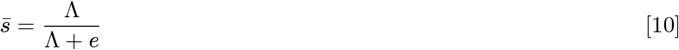

For our model this expression is equivalent to the *fundamental niche* for the specific life history strategy in question and, like Levins expression for metapopulation occupancy, we can describe the carrying capacity or *realized niche* as:

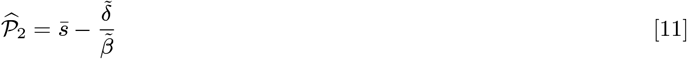

Where, 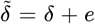. In the case where *e* = 0 and *c* = 0 we return to the canonical equation for metapopulation occupancy (eq. 2).

In the next section we compare the analytic results of the meanfield with numerical simulations of the IPS. We use a von Neumann neighborhood for both propagule production and patch construction. The size of the neighborhood is calculated accordingly:

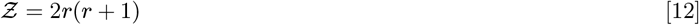

Unless specified otherwise, we set the range of the neighborhood (*r* = 1) i.e. 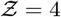 and the lattice size 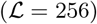. We explored the 2D parameter space generated by life-history trade-off (*c*) and the dynamics of the ephemeral landscape (*e*) with respect to occupancy 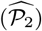 through Monte Carlo methods.

## 2. Results For The Niche Construction Model

### A. Comparison of Mean-Field Approximation and Interacting Particle System Simulations

From figure 2, it is clear that the steady state equilibrium of the MFA overestimates the region of parameter space where the metapopulation can persist. Comparing the plots in figure 2 we see the curve *e*(*c*) defines the population extinction threshold. MFA and IPS alike, the optimal trade-off between propagule production and niche construction is landscape dependent. Tracing along the edge of the x-axis (where the landscape is almost static) we see the highest occupancy where little investment is made towards niche construction. This is due to the small impact of (*e*) on the outcome of the metapopulation. As this rate of habitat destruction increases, the minimalist niche construction strategies die out and a moderate trade-off between the two allows the metapopulation persistence. Notably, the vertex of the extinction threshold curve (the trade-off strategy that persists in the broadest range of *e*) is around *c* = 0.4 for both IPS and MFA, similar to the conclusion made by Krakauer et. al for their single strategy system.

**Fig. 2.**
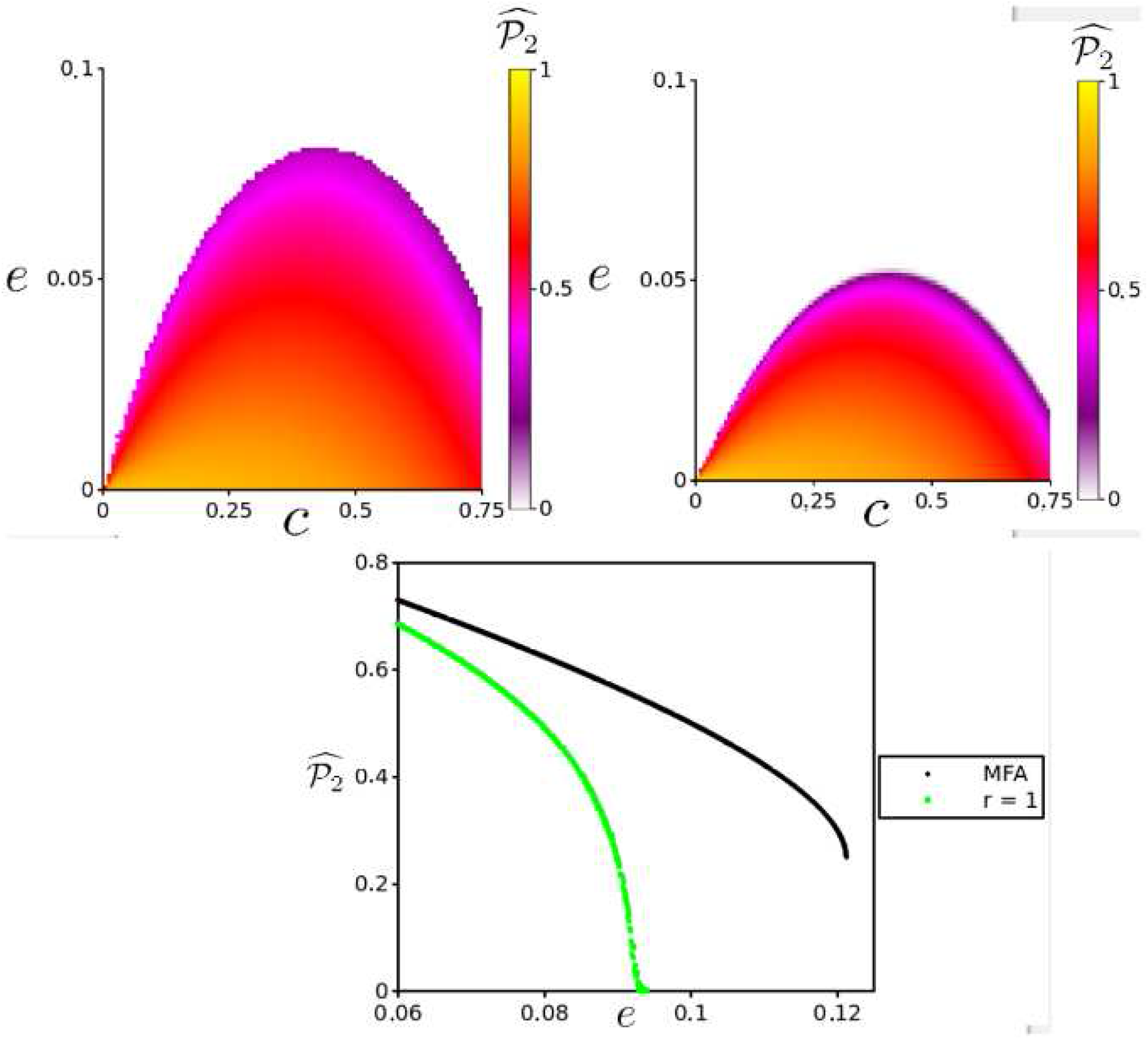
**Top:** A comparison of the MFA (Left) and the IPS (Right) parameter spaces, with the x-axis signifying the life-history strategy adopted (*c*) and the y-axis the hostility of the landscape (*e*). For both *δ* = 0.1 and *α* = 1. For IPS, simulations ran for 5000 times steps, where, upon reaching the invariant measure, a mean occupancy 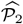 was calculated from the following 250 time steps. In total, 10,000 simulations were run by sweeping the parameter space {*c, e*} creating a 100×100 matrix. **Bottom:** Divergence in the qualitative behavior of the niche construction model. For the MFA a discontinuous transition exists for the order parameter 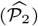, whereas the discontinuity disappears in the SEM for *r_c_* = 1 and we are left with a continuous transition, similar to the contact process. Other parameters for MFA and SEM {*α,δ,c*} = {1, 0, 0.4}.

In figure 3 a simulation shows a successful trade-off from propagule production to niche construction, allowing the population to persist even when confined to small clusters. These clusters are able to sustain themselves and create the corridors within the landscape (See Transects on R.H.S of figure 3), thus escaping extinction. Although individual patches are fleeting, the metapopulation persists. It should be noted that near the extinction threshold, as shown in figure 3, the occupied patches always self-organize into these clusters for *r_c_* = 1. This landscape level spatial organization has been recognized as an early warning signal for arid ecosystems about to transition to a desert state (49).

**Fig. 3.**
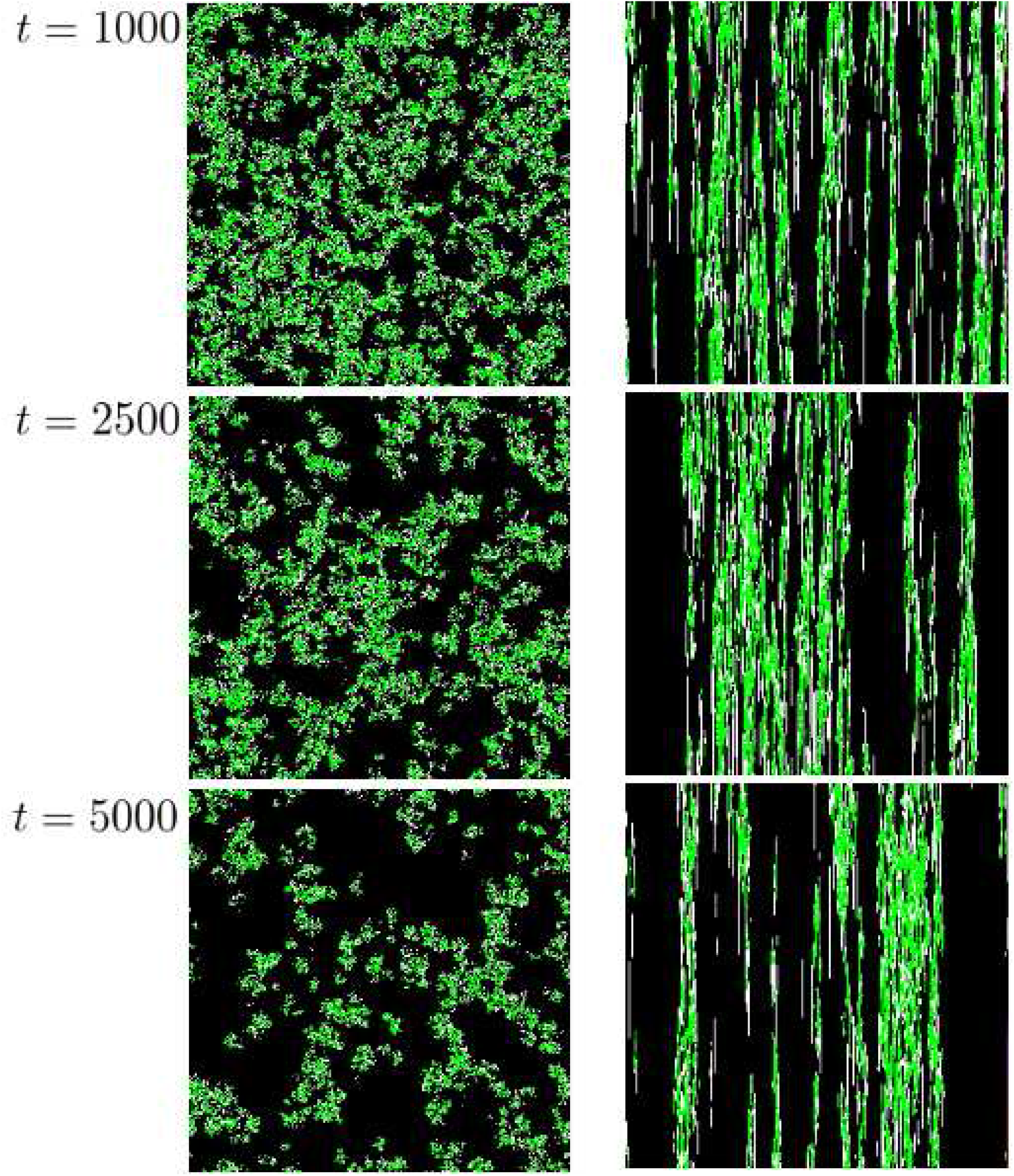
**Left:** Snapshots of the particle system at *t* = 1000, 2500, 5000. Occupied (green), Vacant (white) & Destroyed (black) sites. **Right:** A 1D spatial transect of the particle system for 256 time steps. The final snapshot/transect was taken after the system had reached the invariant measure. Parameter values used for this simulation were at the edge near the extinction threshold (See IPS in Figure 2); {*α, δ, c, e*} = {1, 0.1, 0.25, 0.045}

While the quantitative difference in the MFA and IPS extinction threshold is expected, the qualitative differences observed between MFA and IPS were not anticipated. Following the behavior of the deterministic MFA sudden transition occurs at *e*(*c*)_*crit*_. Considering this result, one would expect the IPS to transition similarly (i.e. discontinuously) to the absorbing state. Instead we are left with a continuous phase transition, much like the one documented for the contact process and more broadly a characteristic of the Directed Percolation Universality Class. Typical of continuous phase transitions, power law behavior of the order parameter 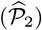 is observed as it approaches the critical value for extinction/habitat destruction (*δ_c_, e_c_*) in both CP and NC particle systems (Figure 4). Furthermore, both display similar critical slowing-down divergence of the relaxation time as shown by simulations run for values below, at and above the critical point.

**Fig. 4.**
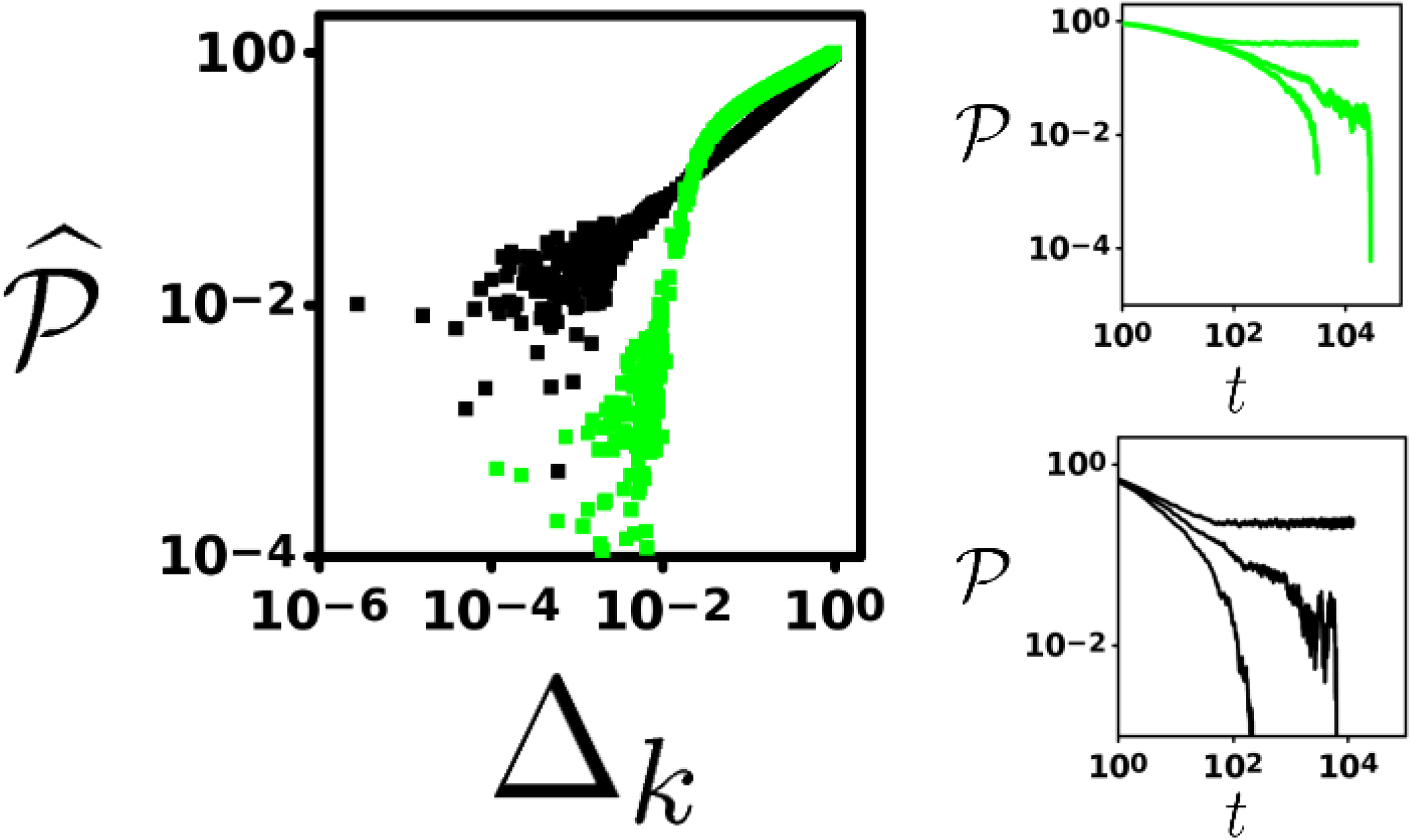
**Left:** Log-log plot showing critical behavior of the order parameter (long-term occupancy, 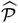) as we approach the critical point *δ_crit_* and *e_crit_* for the contact process (black) and niche construction (green) model, respectively. As these systems near the critical point (Δ_*k*_ ≡ | *k* − *k_crit_* | for *k* = *δ, e*), the order parameters display power law behavior with unique critical exponents (indicated by the unique slope for each). The critical exponent for the contact process determined from this slope (≈ 0.6116) holds up well against previous estimates (46). Parameters for the niche construction model {*α,δ,c*} = {1, 0, 0.4}. **Right:** Dynamical behavior for the Niche Construction model (Top) and Contact process (Bottom) for *k* < *k_crit_* (subcritical), *k* = *k_crit_* (critical) and *k* > *k_crit_* (supercritical), from top to bottom in each plot. Divergence of relaxation time to extinction at the critical point is indicative of continuous phase transitions.

In light of these startling results, we investigated the effect of niche construction neighborhood radius (*r_c_*) on the rate of range expansion by the metapopulation starting from a single occupied site. During the expansion process the metapopulation is both colonizing vacant sites and renewing destroyed sites for future colonization. The lattice quickly becomes divided into two distinct spatial regions, i) A mosaic of occupied, vacant and destroyed sites which maintains its heterogeneity via the extinction (*δ*), destruction (*e*) and construction (*c*) dynamics and ii) A homogeneous vacuum of destroyed sites where no local populations are nearby to combat the global destruction rate (*e*). While the exact configuration within region I of the lattice is constantly changing stochastically, a dynamical internal homeostasis allows the metapopulation to not only persist, but expand as long as the landscape/life-history dynamics are below *e*(*c*)_*crit*_. Interestingly, at the boundary of these two spatial regions we observe an edge effect. It is here, at the edge, where populations along the periphery of the metapopulation mosaic can access the nearest sites of region II, allowing them to convert destroyed sites into vacant sites, facilitating future expansion. As *r_c_* is extended we observe a steady increase in the rate of range expansion eventually leading to the same long-term carrying capacity 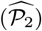 at which point region II vanishes (Figure 5). Intuitively, a larger *r_c_* means a greater portion of occupied sites are able to convert destroyed sites into vacant sites at the edge. Furthermore, periphery occupancy is able to renew destroyed sites beyond adjacent sites within the boundary between regions I and II, therefore widening the gap between regions I and II which in turn increases the chance for propagules to land on vacant sites. As the gap between regions I and II expands this edge becomes ‘fuzzier’ since this expansion is not as tightly coupled as solely adjacent sites (*r_c_* = 1).

**Fig. 5.**
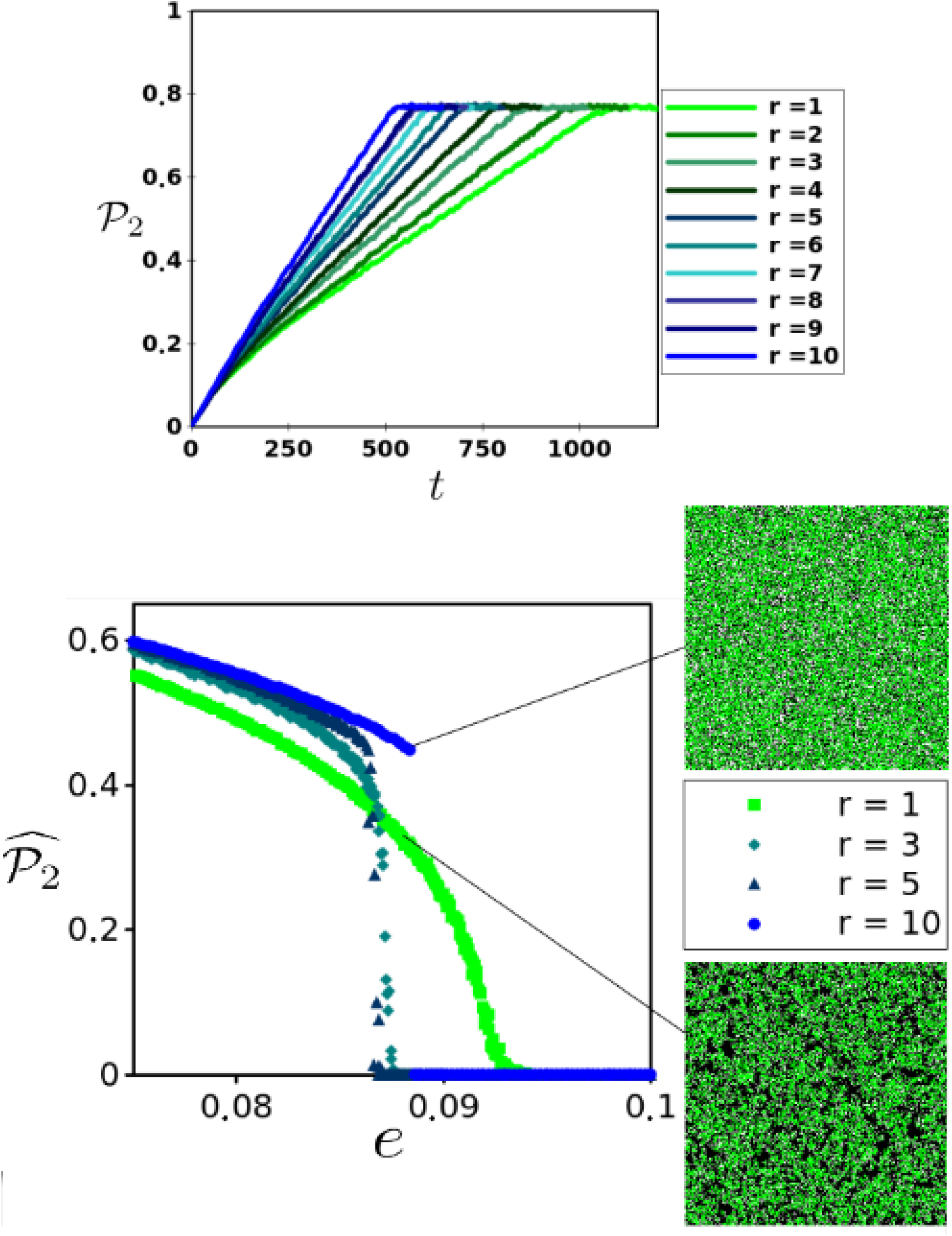
**Top:** Range expansion for niche constructor strategies with different *r_c_*. Identical 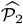 reached for *e* ≪ *e_crit_*. Simulations were run with the following parameter values {*α,δ,c,e*} = {1, 0.1, 0.4, 0.01}. **Bottom:** Long-term behavior for different *r_c_* around *e_crit_* (Left). Parameters used for simulations were {*α,δ,c*} = {1, 0, 0.4}. Snapshots showing differences in spatial clustering taken from simulations at *e* = 0.088 for *r_c_* = 10 and *r_c_* = 1.

While an increase in *r_c_* does not come at an explicit additional cost since we consider the cost fixed for niche construction 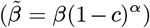, it has surprising implications for the resiliency of the metapopulation. As we approach *e*(*c*)_*crit*_ from below, 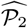 begins to increase steadily with larger *r_c_*. However, we observe a qualitative change in the behavior of the model at *e*(*c*)_*crit*_ for *r_c_* > 1; as *r_c_* increases the transition to the absorbing state becomes steeper, eventually displaying a discontinuity, as shown for *r_c_* = 10, and similar to the MFA. Additionally, the spatial clustering exhibited for *r_c_* = 1 near *e*(*c*)_*crit*_ disappears. This last observation happens to be the key to understanding why the extinction threshold changes qualitatively as *r_c_* increases. To do so, it is useful to return to the idea of 2 spatial regions which we discussed when considering range expansion starting from an initial cluster of occupied sites in a sea of destroyed sites. Now consider starting with a fully occupied lattice 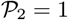. For small *r_c_* as *e* → *e_crit_* the metapopulation structure becomes fragmented, but clusters continue to persist (Figure 3). These clusters of occupied sites (Region I) drift through the sea of uninhabitable sites (Region II) occasionally fusing with other clusters and occasionally going extinct. Since this drift depends on their ability to renew adjacent sites at the edge they benefit from the highly condensed internal structure of the local cluster and the tight coupling with recent niche construction events. Internally, they maintain homeostasis while capitalizing most effectively on renewed sites at the edge because any renewed site is likely adjacent to at least one viable population. If *e* = *e_crit_* then the relaxation time to extinction behaves as a power law as expected for continuous absorbing state phase transitions (46). As *r_c_* increases, clustering is lost for the same reason it aids in range expansion; instead of the “fuzziness” being generated at the edge of the expanding metapopulation it exists over the entire lattice. Therefore, when *e* = *e_crit_*, given enough time an area of the lattice will return to Region II. In contrast to the sub-critical regime, Region II now expands, fragmenting the metapopulation. Since the the niche construction neighborhood is large, no clustering occurs and the system experiences a rapid relaxation time to the absorbing state.

## 3. Addition to the Model

### A. Competing for Niche Space

To further explore the colonization-niche construction trade-off, we introduce an additional particle strategy to see how competition between two life-history strategies affects the long-term outcome of metapopulation persistence in this ephemeral landscape.

We consider the basic contact process (See *Section 2.1.*) as the alternative strategy. Since this strategy does not partition effort to local patch renewal it has a higher 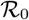 than the niche constructor. Given a static landscape with two competing particle strategies, the larger 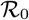 will push the smaller 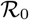 strategy to eventual extinction. This example of the competitive exclusion principle has been proven rigorously (50) in the context of multiple competing contact processes. However, any non-zero value for *e* requires some investment *c* within the metapopulation. Therefore we would expect the contact process to behave like a fugitive strategy (51), more capable of colonizing vacant sites, but unable to maintain stable concentrated patch clusters through time. For this model we do not assume any competitive hierarchy between the two strategies. An updated schematic of the complete model is provided (Figure 6).

**Fig. 6.**
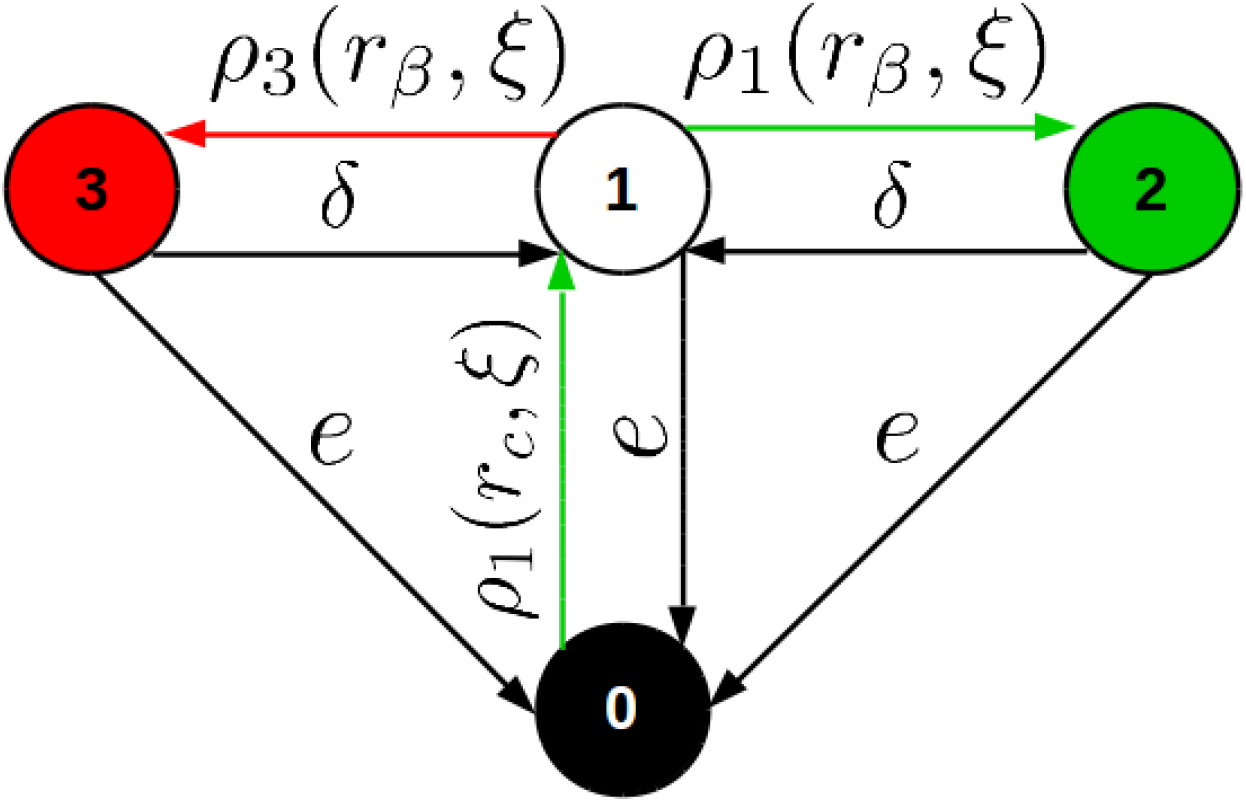
The only adjustment from the previous model is the introduction of the contact process (red) as an alternative strategy to the niche constructor(green). Sites in the lattice now acquire one of 4 possible states, 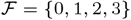.

## 4. Results For The Competition Model

We first compare the dynamics of the model when the constructed niche is completely uncoupled to the NC (i.e. the neighborhood equals the entire lattice), versus when it is tightly coupled to the NC strategy (*r_c_* = 1) in Figure 7. We see that for the global case, the CP strategy out-competes the NC strategy globally, leading to its extinction. Once this occurs, nothing prevents the lattice from becoming completely destroyed. For future reference, we refer to this outcome as *Ecological Suicide* since we do not consider evolving populations. In the case where *r_c_* = 1 we see a drastically different outcome, in which both strategies coexist.

**Fig. 7.**
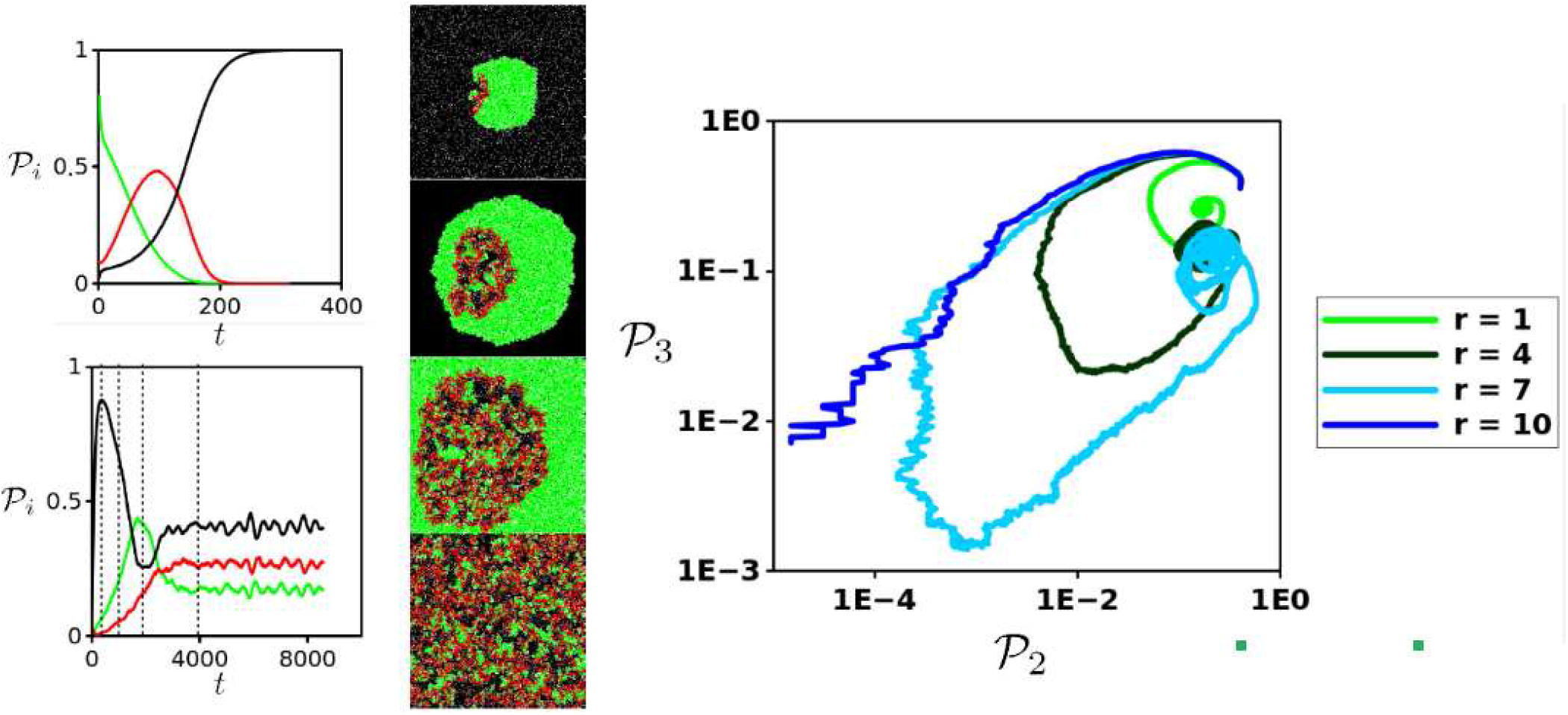
**Left:** Population dynamics for global (Top) and *r_c_* = 1 (Bottom) niche construction neighborhood following niche constructor (green) contact process (red) strategies along with destroyed habitat (black) in the coexistence regime: {*α,δ,c,e*} = {1, 0.1, 0.4, 0.01}. Four snapshots taken at *t* = 250, 1000, 2000, 4000 from top to bottom, temporal locations of snapshots are indicated with vertical dashed lines. Frame 1 shows the CP population being fragmented from vacant sites due to habitat destruction, but rescued by the NC population expanding via the construction of viable habitat. Once the CP population establishes itself within the confines of the surrounding NC particles (frame 2 and 3) they can begin exploiting this renewed habitat and freshly vacated sites (generated by *c* and *δ*, respectively) due to their superior colonization rate 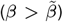. Notice the trail of destroyed habitat as this CP population follows the leading wave of the NC population outward, occasionally becoming fragmented from empty sites and therefore destined for local extinction. Eventually, the system equilibrates, though small fluctuations continue due to the local oscillations and the finite size of the landscape, see (52) for a discussion on this with respect to lattice models. **Right:** Trajectories of the NC-CP phase space for different *r_c_* values. Larger values of *r_c_* increase the likelihood of stochastic extinction during the transient period. Oscillations are caused by range expansion/exploitation of the NC/CP strategies. If the NC metapopulation can escape the first wave of exploitation from the CP strategy then coexistence is likely to occur.

Coexistence is possible because NC sites have an advantage in adjacency to the constructed niche, while CP sites have a colonization advantage, this adjacency advantage diminishes for increased *r_c_*. CP sites chase clusters of NC sites leading to the oscillations observed in the bottom plot for Figure 7. With increased *r_c_*, amplitude increases because this more easily facilitates the CP strategy from exploiting renewed sites generated by NC populations. While increased *r_c_* leads to a faster rate of range expansion for the single strategy model, this benefit is lost in the competition model for the reasons discussed.

If NC populations become surrounded by CP sites they are likely to go extinct, unless a gap is created by either death or destruction events. Therefore, the NC strategy is only able to persist if they can access the interface with region II, as discussed in the previous section.

Along with the previously observed instances where the dynamics of the landscape push the NC metapopulation to extinction, it is clear, 3 regimes of the parameter space appear from these simulations (Figure 8), described as follows: i) CP goes extinct due to fragmentation in the dynamic landscape with the NC strategy able to persist at the same 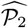 as in the single strategy model ii) coexistence between NC and CP where the CP is the exploiter strategy, persisting in the vacancies made by local extinction and niche construction of the NC strategy leading to an overall lower steady state equilibrium for occupied sites, and iii) *Ecological Suicide*; where CP over-exploits NC, forcing NC to extinction by sufficiently out-colonizing niche constructed sites to the point of enclosing small clusters of NC populations. If the NC strategy cannot escape the CP it eventually becomes extinct resulting in the subsequent extinction of the remaining CP population. As for the coexistence regime (ii), we see a steady rise in CP occupancy at the expense of NC occupancy as NC invest more towards niche construction (*c* → 1). The more NC invests in niche construction the easier it is for CP to exploit because the constructed niche is less tightly coupled to the NC populations.

**Fig. 8.**
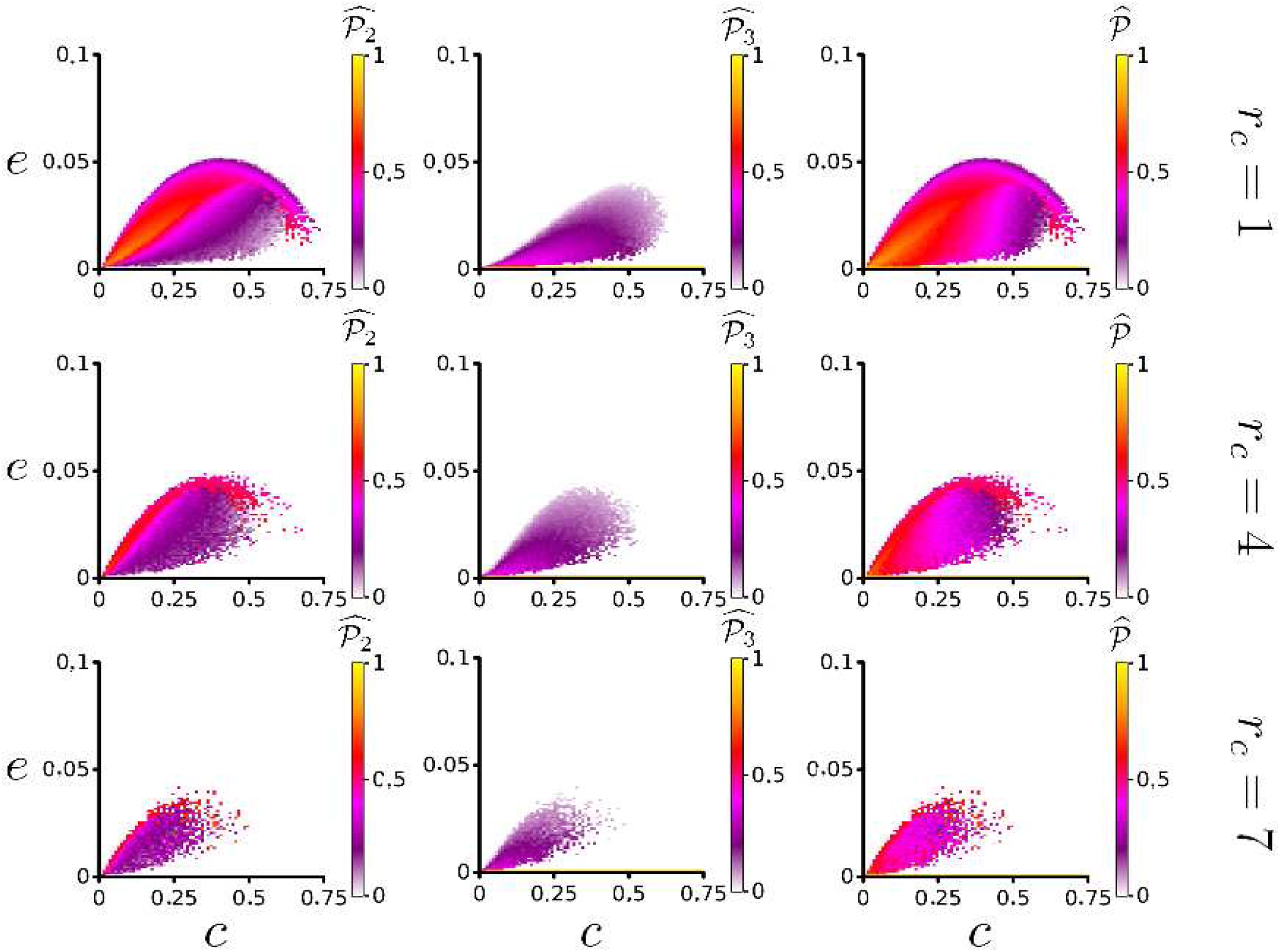
Parameter space {*c, e*} for the model with competition. Left, center and right columns correspond to mean occupancy for NC, CP and both strategies 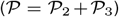, respectively. Three niche construction neighborhoods *r_c_* = 1,4, 7 from top to bottom. {*α,δ*} = {1, 0.1}. As *r_c_* increases ecological extinction expands where CP overexploits NC. The noisiness of this transition, exemplified by spotted pattern of the parameter space, is explained in the main text.

Interestingly, for *r_c_* > 1, stochastic events on the lattice can determine the long-term outcome of the community. When a simulation starts, CP sites quickly colonize territory around NC sites which self organize into clusters. For *c* ≫ 0 clusters of NC become encircled by CP sites, which eventually choke-out NC populations leading to ecological suicide. If however, they are able to slip past CP sites and reach region II of the lattice, they can continue to expand their range, at which point CP sites connected by vacancy will continue to follow their expansion. Alternatively, if CP sites become separated entirely from NC sites, CP populations will stay fragmented from vacant sites and eventually be vanquished by this self-inflicted fragmentation and the NC metapopulation will return to the long-term behavior of regime I. It is these mechanisms which lead to the noisy behavior of the model at the edge of the extinction threshold, and consequently, this noisiness is increased as *r_c_* is increased.

## 5. Discussion

This model is similar in structure to one previously described, aimed at understanding the impact of facilitation (*f*) by vegetation in a dynamic landscape of viable and degraded patches (53). It is clear, the scope of our models vary which in turn has subtle effects on the dynamics of the systems studied, for example, their consideration of only vacant patches as susceptible to degradation. This is a logical assumption for their model as arid vegetation is well adapted to retain moisture in such extreme climates. Surprisingly, these subtle changes lead to strikingly different outcomes at the extinction threshold. In particular, they observed both continuous and discontinuous transitions in their lattice model and pair approximation for nearest neighbor facilitation, *r_f_* = 1, where the outcome depended on other life-history and environmental parameters. In addition, while their work identified similar spatial clustering of vegetation for both continuous and discontinuous transitions, we see only the continuous transition displaying such self-organizing behavior at the scale of the landscape. In fact, it is this discrepancy caused by a decoupling of the constructed niche to the metapopulation which we suspect leads to the discontinuity in our model. On this note, while this previous work did not consider *r_f_* larger than 1, our results suggest that this clustering behavior may disappear if the range for facilitation is only slightly larger than immediate adjacency. This is significant because clustering is widely described as an important early warning signal for critical transitions in arid ecosystems. Determining the true universality of this pattern formation behavior is an ongoing line of investigation, and one which will feedback into our ability to predict and prevent environmental catastrophes (54).

Our goal here has been to implement a spatial instantiation of the constructed niche as defined by Krakauer et al. (20), through the interacting particle system framework, in order to determine if a spatial perspective would reveal new and interesting insights into the ecological consequences of niche construction in ephemeral landscapes. While some observations made by Krakauer et al. were reconfirmed here in the spatial model, for example the optimal trade-off between construction and colonization in the single strategy system, new significant findings were also uncovered; for instance, in cases where niche construction is spatially local (*r_c_* = 1) leading to the emergence of clustering. This clustering in turn allowed the metapopulation to maintain itself where larger *r_c_* could not. This outcome was reminiscent of the bi-stability predicted by the MFA. In contrast to these mean field predictions, no such behavior was observed for the spatial model when *r_c_* was limited to adjacent sites. Applying finite-size scaling theory through Monte Carlo simulations revealed a continuous phase transition which was steeper than, but qualitatively similar to the Contact Process. As we increased *r_c_* for the IPS this transition became more drastic, until it eventually vanished and the discontinuity predicted in the MFA was recovered.

Continuing our study of the single strategy model, we investigated the impact of *r_c_* on range expansion. As expected, increased niche construction neighborhoods led to an increased rate of range expansion. Therefore, before implementing a second strategy into the model a trade-off existed for the niche construction strategy between the rate of range expansion and susceptibility of the metapopulation to critical transitions.

In light of the trade-off observed for the single strategy model, it is interesting to consider how the direction and strength of selection on both the proportion of niche construction (*c*) and the range (*r_c_*) would play out for a system where population strategy varied. We envision range expansion experiments implemented in microbial systems to be an ideal framework for testing these predictions (55).

Next, we studied the two strategy model, where the regular contact process behaved as an exploiter strategy to the niche constructor. As expected, increased niche construction range allowed for greater exploitation. For regions of the parameter space, the system experienced what we referred to as ecological suicide, where CP pushed NC to extinction, thus sowing the seeds to their own demise. Unexpectedly, extinction due to ecological suicide was not as clearly defined in our parameter space as the single strategy phase transitions. For example, unlike in the single strategy model where we located *e_crit_* through finite scaling methods, we are unable to define critical neighborhood size where ecological suicide begins. This is possibly due to the stochastic nature of its mechanisms; wherein, clusters of NC sites capable of escaping local extinction via competition are freed from the exploiter strategy allowing them to reach equilibrium values equivalent to the single strategy model. For *r_c_* = 1, ecological suicide is most probable when *e* is small and *c* large because the NC reproductive ratio is compromised by a high investment in niche construction. This allows the CP to quickly out-compete and surround NC before being fragmented by patches of destruction. As *r_c_* increases the speed with which the CP strategy is able to surround NC increases. The expanding area of the parameter space where ecological suicide occurs as *r_c_* increases exemplifies the tragedy of the commons described by (20).

This result emphasizes the importance of spatial structure on the long-term and global outcome of niche constructor/exploiter metapopulation dynamics. Notably, how a few stochastic events changing the connectivity of patches between populations of niche constructing and exploiting strategies lead to drastically different outcomes. Specifically, the initial conditions of these founder populations (clusters of occupied sites surrounded by destroyed sites) can exhibit both stochastic (dominated by random chance events) and deterministic phases (predicted by the proportion of NC vs. CP sites in a cluster) (56). In this case, the transition between these two phases is determined by *r_c_.*

As a point of departure, we considered space as the only dimension of the niche by which a population can construct upon. Adding other resources or phenotypes would enrich the behavior of the model. For example, the addition of combinatoric niche construction reactions possibly similar in structure to Fontana’s Algorithmic Chemistry (57), where now different varying phenotypes can lead to expansion into adjacent possible areas of the niche space. It would be interesting to see how the spatial heterogeneity of these resources may lead to further exploitation or cooperation behavior and how the position of these resources along the excludability spectrum plays a role in the predictability of long-term outcomes.

## 6. Conclusion

We envision this trade-off to be a generalized predicament faced by populations and whose solutions to this dilemma mark the most significant transitions in the evolution of life on earth (18). Of particular interest is the origins of multicellularity, which was most likely achieved through the division of labor between maintenance and replication (4, 5). Here we developed a ‘toy ecology’ within the interacting particle system framework which we believe reproduces some of the conceptual dilemmas faced at this transition from population to individual. Furthermore, these findings have implications for the study of ecosystems stability and susceptibility to critical transitions. Understanding which real world ecosystems are likely to behave as continuous vs. discontinuous phase transitions is crucial for their conservation and management.

### A. Mean Field Equilibrium For The Niche Construction Model

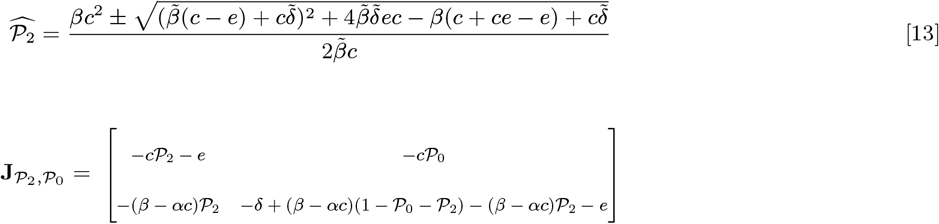

The eigenvalues all have real parts that are negative therefore the system is stable near the stationary point.

## ACKNOWLEDGMENTS

We thank Alfredo L’Homme and Janneke Noorlag for valuable discussions and providing feedback on earlier versions of this manuscript. We thank financial support from FONDECYT 1150430, Conicyt, Chile. MTW also thanks financial support from VRI for their Instructor Scholarship.

